# TOP-2 is differentially required for the proper maintenance of cohesin on meiotic chromosomes in *C. elegans* spermatogenesis and oogenesis

**DOI:** 10.1101/2021.12.26.474225

**Authors:** Christine Rourke, Aimee Jaramillo-Lambert

## Abstract

During meiotic prophase I, accurate segregation of homologous chromosomes requires the establishment of a chromosomes with a meiosis-specific architecture. Sister chromatid cohesins and the enzyme Topoisomerase II are important components of meiotic chromosome axes, but the relationship of these proteins in the context of meiotic chromosome segregation is poorly defined. Here, we analyzed the role of Topoisomerase II (TOP-2) in the timely release of sister chromatid cohesins during spermatogenesis and oogenesis of *Caenorhabditis elegans*. We show that there is a different requirement for TOP-2 in meiosis of spermatogenesis and oogenesis. The loss-of-function mutation *top-2(it7)* results in premature REC-8 removal in spermatogenesis, but not oogenesis. This is due to a failure to maintain the HORMA-domain proteins HTP-1 and HTP-2 (HTP-1/2) on chromosome axes at diakinesis and mislocalization of the downstream components that control sister chromatid cohesion release including Aurora B kinase. In oogenesis, *top-2(it7)* causes a delay in the localization of Aurora B to oocyte chromosomes but can be rescued through premature activation of the maturation promoting factor via knock-down of the inhibitor kinase WEE-1.3. The delay in Aurora B localization is associated with an increase in the length of diakinesis chromosomes and *wee-1.3* RNAi mediated rescue of Auorora B localization in *top-2(it7)* is associated with a decrease in chromosome length. Our results imply that the sex-specific effects of Topoisomerase II on sister chromatid cohesion release are due to differences in the temporal regulation of meiosis and chromosome structure in late prophase I in spermatogenesis and oogenesis.

## INTRODUCTION

Accurate segregation of chromosomes to daughter cells during mitosis and meiosis is essential for the maintenance of genomic stability. Sister chromatid cohesion (SCC) plays a critical role in this process by ensuring sister chromatids are held together until their regulated separation at anaphase. SCC is mediated by a tripartite protein ring complex that topologically entraps sister chromatids (Nasmyth & Haering, 2009). The cohesin core complex consists of four subunits: two structural maintenance of chromosomes (Smc1 and Smc3) proteins, a Kleisin subunit (Rad21/Scc1), and an accessory protein (Scc3/SA1/SA2) (Rankin, 2015). In mitosis sister chromatids remain tethered together until the transition from metaphase to anaphase. At the onset of anaphase Rad21 is cleaved by the protease separase, which allows separation of the sister chromatids to opposite poles of the spindle. The release of SCC must be tightly regulated to prevent the precocious separation of sister chromatids. Precocious separation of sister chromatids leads to severe defects in chromosome segregation, which has been associated with tumorigenesis and developmental disorders (Barber et al., 2008; Krantz et al., 2004; Tonkin et al., 2004; Vega et al., 2005).

During meiosis, haploid gametes are generated from diploid cells that undergo a single round of DNA replication followed by two successive chromosome segregation events. SCC establishment and removal during this specialized cell division is even more complex as the accurate segregation of homologous chromosomes at the first meiotic division depends on SCC in the context of crossovers (CO). The combination of SCC and COs ensures the correct orientation of homologous chromosomes on the meiotic spindle (Petronczki et al., 2003). Precocious loss of SCC in oocytes contributes to aneuploidy, a leading cause of miscarriages and birth defects in humans (Hassold & Hunt, 2001).

In meiosis, the kleisin subunit of the cohesin complex is replaced by a meiosis-specific subunit, Rec8 (Rankin, 2015). SCC is released in a stepwise manner during meiosis. First, cohesin is removed at the chromosome arms through phosphorylation of Rec8 by the Aurora B kinase and subsequent cleavage of Rec8 by separase. Protection of Rec8 at the centromeres is mediated by protein phosphatase 2A allowing for the separation of homologous chromosomes at meiosis I while keeping the sister chromatids tethered until meiosis II (Ishiguro et al., 2010; Kitajima et al., 2006; Riedel et al., 2006). During meiosis II, the remaining cohesin complexes at the centromeres are cleaved by separase promoting the segregation of sister chromatids.

Meiotic chromosome structure is also of critical importance for the accurate segregation of homologous chromosomes. Chromosome axes conjoin linear looped arrays of sister chromatids at the loop base. Chromosome axes are composed of cohesin complexes as well as meiosis specific proteins such as the HORMA-domain proteins (Blat et al., 2002; Kim et al., 2014). These axis proteins are required for the assembly of the synaptonemal complex to establish homologous chromosome pairing and meiotic recombination. Also associated with meiotic chromosome axes is the enzyme Topoisomerase II (Gomez et al., 2014; Jaramillo-Lambert et al., 2016; Kleckner et al., 2013; Klein et al., 1992; Mengoli et al., 2014). In mitosis, Topoisomerase II along with SMC proteins and high mobility group (HMG) proteins make up the protein core around which chromosomes are organized (Zickler & Kleckner, 1999). Although Topoisomerase II is part of the chromosome axes of meiotic chromosomes, the relationship between meiotic chromosome structure, establishment and disassembly of SCC, and Topoisomerase II is poorly understood.

In this work, we study the sex-specific connection between SCC release and Topoisomerase II. We find that Topoisomerase II is differentially required in oogenesis and spermatogenesis to maintain SCC until chromosome segregation at anaphase I. When Topoisomerase II is disrupted during spermatogenesis, REC-8 cohesin is precociously removed. This is due to a failure to maintain the HORMA-domain proteins HTP-1 and HTP-2 (HTP-1/2) on chromosome axes after diakinesis. The disruption in HTP-1/2 localization leads to the mislocalization of downstream components that control SCC release including Aurora B kinase. In contrast to spermatogenesis, loss of Topoisomerase II function during oogenesis neither disrupts HTP-1/2 localization nor most downstream SCC release factors. However, localization of Aurora B to oocyte diakinesis chromosomes is delayed in a Topoisomerase II loss-of-function mutant. We propose that the sex-specific effects of Topoisomerase II on SCC release are due to differences in the temporal regulation of meiosis and chromosome structure in late prophase I in spermatogenesis and oogenesis.

## MATERIALS AND METHODS

### Strains

The following strains were maintained at 20°C under standard conditions, N2: wild-type Bristol strain, CB1489: *him-8(e1489)* IV, DG4915: *his-72(uge30)* III*; fog-2(oz40)* V, CV6: *lab-1(tm1791)/hT2*, and ATGSi23: *fqSi23* II *[Prec-8 rec-8 ::GFP 3’UTR rec-8; cb-unc-119(+)]; rec-8(ok978)* IV (Ferrandiz et al., 2018). The remaining strains were maintained at 15°C under standard conditions and are as follows: AG259: *top-2(it7)* II*; him- 8(e1489)* IV (Jaramillo-Lambert et al., 2016), AJL1: *top-2(ude1)* [*it7* recreate] II; *fqSi23 II* [*Prec-8 rec-8 ::GFP 3’UTR rec-8; cb-unc-119(+)*].

### Immunofluorescence and DAPI staining

Hermaphrodites and males were synchronized by picking L4s and incubating at 15°C for 16-24 h. Adult worms were then placed into the 24°C-incubator for 4 hours. Gonads were dissected in 30 μl of 1x Egg buffer (118 mM NaCl, 48 mM KCl, 2 mM CaCl2 · 2H2O, 2 mM MgCl2 · 6H2O, 25 mM HEPES, pH 7.4) and 0.1% Tween 20 on a glass coverslip. After dissection, 15 μl of liquid was removed and replaced with 15 μl 2% paraformaldehyde solution (16% paraformaldehyde from EM Sciences in 1x Egg buffer, 0.1% Tween 20). 15 μl of liquid was removed and a Superfrost Plus microscope slide (Fisher, cat #12-550-15) was placed on top of the cover slip containing the dissected gonads. The slide and coverslip were flipped right-side up, allowed to fix for 5 min then placed into liquid nitrogen. Once frozen, the coverslip was quickly removed, and the slide was immediately submerged in -20°C methanol for 1 min. The slide was washed three times (5 min each) in PBST (1x PBS, 0.1% Tween 20). Following the three washes, the slides were incubated at room temperature for 1 h in a blocking solution (0.7% BSA in PBST). 50 μl of primary antibody diluted in blocking solution was placed on top of the gonads, covered with a parafilm coverslip and incubated overnight (16-24 h) at room temperature in a humidified chamber. Following primary antibody incubation, slides were washed three times in PBST for 5 min each and incubated at room temperature with 50 μl of secondary antibody diluted in blocking solution in a humidified chamber for 2 h. Slides were washed 3 times for 5 min each in PBST, then stained with DAPI (2μg/mL) for 5 min. Slides were washed for a final 5 min in PBST and mounted with Vectashield and a glass coverslip.

For experiments where DAPI staining alone was used, the worms were dissected, fixed, and frozen in liquid nitrogen as above. Once frozen, the coverslip was quickly removed, and the slide was immediately submerged in -20°C methanol for 1 min. The slide was washed one time (5 min) in PBST, removed and incubated with DAPI (2μg/mL) for 5 min in the dark. Slides were then washed for a 5 min in PBST and mounted with Vectashield and a glass coverslip.

Primary antibodies used in this study were: rabbit anti-COH-3/4 (1:4,000) (Severson & Meyer, 2014), rabbit anti-REC-8 (1:100) (Pasierbek et al., 2001), rabbit anti-HTP-1/2 (Martinez-Perez et al., 2008), guinea pig anti-HTP-3 (1:750) (MacQueen et al., 2005), rabbit anti-LAB-1 (1:300) (de Carvalho et al., 2008), rabbit anti-AIR-2 [(1:100) Figure 2A and Figure 6] (Schumacher et al., 1998), rabbit anti-AIR-2 [(1:500) Figure 2B] (Cell Signaling Technology, 2914S), Histone H3 phospho-T3 (1:750) (Millipore, 07-424), and rabbit anti-GFP (1:500) (Novus Biologicals, NB600308). Secondary antibodies used were goat anti-rabbit Alexa Flour-568 and goat anti-guinea pig Alexa Flour-488 (1:200) (Invitrogen, Thermo Fisher Scientific, A11036 and A11073).

### Image collection, processing, and measurements

Z-stack images of dissected gonads were obtained using a Zeiss LSM880 confocal microscope (Carl Zeiss, Inc., Gottingen, Germany) using a 63x objective and Zen software. For each gonad imaged, a full range of focal planes were selected to include the complete gonad, with a constant Z-step of 0.2 μm. Image processing and analysis was done using Fiji Is Just ImageJ [Fiji, (Schindelin et al., 2012)]. The range of focal planes chosen for z-stack projections in figures was chosen depending on the image and whether the image was of a male or a hermaphrodite gonad. Images were taken using identical imaging parameters, but brightness and contrast were adjusted to allow for better visualization. Images used for chromosome length measurements were deconvolved using Huygens Profession 20.10 software. The deconvolved images were visualized in 3D using Imaris x64 9.7.2 software and measurements were made by visually delimiting individual diakinesis bivalents and each bivalent length plotted using Prism (GraphPad Software).

### RNAi

RNA was introduced into worms using the feeding method of (Timmons et al., 2001). RNAi clones [*smd-1* (no phenotype negative control) and *wee-1.3*] were obtained from either the Ahringer library or the OpenBiosytems library. Three to five L4 hermaphrodites were placed on a single 35 mm MYOB plate spotted with 50 μl of *E. coli* HT115 [DE3] containing plasmids for RNAi knockdown. Hermaphrodites were allowed to lay embryos and 5 days post-hatching (15°C) ∼20 L4 F1 progeny were transferred to fresh RNAi plates. After 16-24 hours the F1 hermaphrodites were transferred to 24°C for 4h. After incubating at 24°C, worms were transferred to a glass coverslip for gonad dissection and immunostaining.

### Embryonic viability mating assay

Single L4 males of the indicated genotype (**Figure S4E**) were mated with a single L4 *fog-2(oz40)* female at 20°C on a MYOB plate spotted with OP50 *E. coli*. Every 24 h the adult worms were transferred to fresh plates. This was repeated twice and on the third day the adult male and female were discarded. The number of live progeny and dead embryos were counted for each 24 h period and the percent embryonic viability was calculated. Three independent trials were performed for each mating assay. N2 male crosses N=20 and *lab-1(tm1791)* N=22.

### CRISPR/Cas9-mediated genome editing

CRISPR-mediated genome editing to recreate the *top-2(it7)* R828C mutation in *fqSi23* [*prec-8; rec-8::gfp; rec-8 3’UTR; cb-unc-119(+)*] was done as in (Jaramillo-Lambert et al., 2016). A single line was generated carrying the R828C mutation in *fqSi23* and given the allele designation *ude1*.

## DATA AVAILABILITY

Strains are available upon request. The authors state that all data necessary for confirming the conclusions are presented in the article and figures. Figure S1 shows the localization of COH-3/4 in spermatogenesis. Figure S2 shows the dynamics of HTP-3 and H3pT3 localization in spermatogenesis. Figure S3. Shows HTP-1/2 localization pachytene of spermatogenesis. Figure S4 shows LAB-1 localization dynamics in spermatogenesis. Supplementary material is available online at FigShare.

## RESULTS

### TOP-2 is required to prevent the premature loss of REC-8 cohesin in the male germline

Previous work from our laboratory showed that the enzyme Topoisomerase II, TOP-2, is differentially required for homologous chromosome segregation during meiosis I in male and female germlines (Jaramillo-Lambert et al., 2016). Analysis of *top-2(it7)*, a temperature sensitive allele, in both male spermatogenesis and hermaphrodite oogenesis showed that at the nonpermissive temperature (24°C) segregation of homologous chromosomes is disrupted during the first meiotic division of spermatogenesis, but the meiotic divisions of oogenesis are not affected (Jaramillo-Lambert et al., 2016). However, the molecular mechanisms requiring differences in TOP-2 localization and/or activity at play in the two sexes is not known. In other organisms Topoisomerase II has been implicated in chromosome structure and condensation (Gomez et al., 2014; Hartsuiker et al., 1998; Li et al., 2013). We previously demonstrated through a time course analysis that TOP-2 functions late in meiotic prophase I of *C. elegans* (Jaramillo-Lambert et al., 2016) around the time of major chromosome remodeling as the chromosomes prepare for segregation. To determine whether TOP-2 is required for chromosome remodeling and structural maintenance during late meiotic prophase, we examined the localization patterns of chromosome structural components in control [*him-8(e1489)*] and *top-2(it7); him- 8(e1489)* male germlines at the nonpermissive temperature (24°C). The *him-8(e1489)* allele (**H**igh **I**ncidence of **M**ales) produces ∼30% male progeny due to chromosome nondisjunction facilitating assessment in males (Hodgkin et al., 1979; Phillips et al., 2005). As previous work in a mammalian system found a link between Rec8 (chromosome structural component) removal and Topoisomerase II alpha decatenation activity (Gomez et al., 2014), we first examined the localization of the *C. elegans* meiosis-specific cohesin, REC-8. In control males, REC-8 localized to the long arms of the bivalents at metaphase I through anaphase I and remained associated with sister chromatids until anaphase II (**Figure 1A & B**). In *top-2(it7); him-8(e1489)* spermatogenic germlines we found that REC-8 localized to the chromosomes through metaphase I but was absent from the aberrant chromosome structures in anaphase I (**Figure 1A & B**). As previously demonstrated, the chromosomes in *top-2(it7)* fail to proceed past anaphase I and no metaphase II or anaphase II nuclei were observed. This data indicates that REC-8 is prematurely removed or lost from chromosomes during spermatogenesis of *top-2(it7)*.

**Figure 1.**
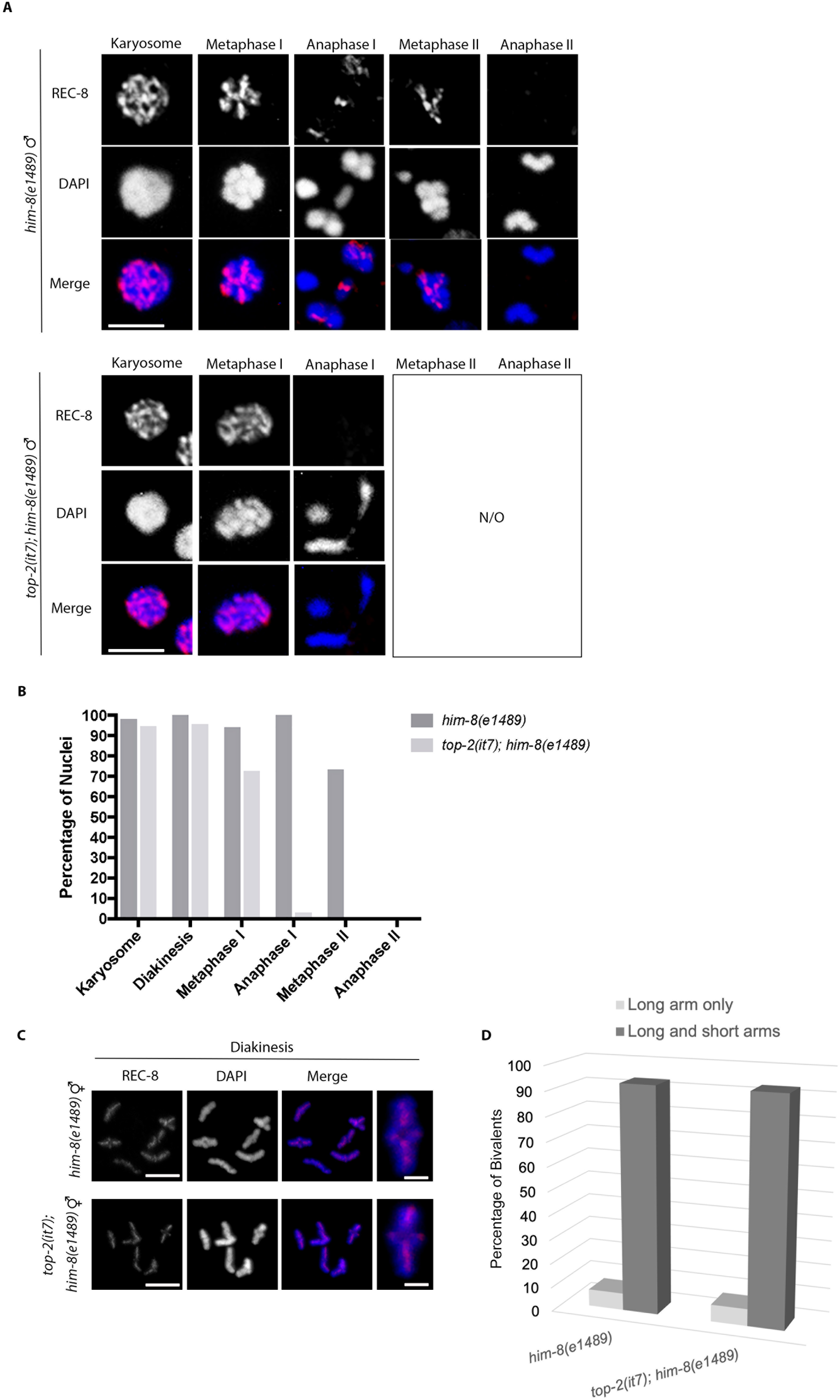
REC-8 is prematurely removed from anaphase I meiotic chromosomes in *top-2(it7)* spermatogenic germlines but not oogenesis. Immunostaining of REC-8 (red) in control [*him-8(e1489)*] and *top-2(it7); him-8(e1489)* during spermatogenesis (A) and oogenesis (C) counterstained with DAPI (blue). (A) Images are Z-projections of nuclei from the karyosome through anaphase II. (B) Quantification of nuclei positive for REC-8 staining in spermatogenesis. (C) Z-projections of diakinesis nuclei during oogenesis (-1 oocyte). Right panels are single magnified bivalents. REC-8 localizes properly in *top-2(it7); him-8(e1489)* hermaphrodites to both arms of the diakinesis bivalent. (D) Quantification of bivalents from the -1 oocyte with REC-8 staining on both the long arm and short arm of the bivalent or the long arm only. N/O = not observed. Scale bar = 5 μm. Magnified bivalent scale bar = 2 μm.

In diakinesis oocytes REC-8 is found on both the long and short arm of the bivalents until late diakinesis or metaphase I when REC-8 is removed from the short arm (de Carvalho et al., 2008; Ferrandiz et al., 2018; Harper et al., 2011; Rogers et al., 2002; Severson & Meyer, 2014). We confirmed that REC-8 localizes to both arms of the diakinesis bivalents in control *him-8(e1489)* oogenesis (**Figure 1C & D**). In *top-2(it7); him-8(e1489)* oogenesis REC-8 localized to both arms of the diakinesis bivalents as in control oogenesis (**Figure 1C & D**). This contrasted with the premature removal of REC-8 in *top-2(it7)* spermatogenesis consistent with the sperm-specific segregation defects of *top-2(it7)*.

Along with REC-8, *C. elegans* has two additional meiosis-specific cohesins, COH-3 and COH-4. COH-3 and COH-4 are 84% identical and are referred to as one-unit COH-3/4 (Severson et al., 2009). During prometaphase I of oogenesis, COH-3/4 and REC-8 occupy different chromosomal domains of the bivalent (Severson & Meyer, 2014; Woglar et al., 2020). COH-3/4 localizes to the short arms of the bivalent and REC-8 first localizes to both the short and long arms and then is restricted to the long arms (Severson & Meyer, 2014). Examination of the meiotic divisions in spermatogenesis of *top-2(it7); him-8(e1489)* germlines (24°C) found that COH-3/4 localize as foci in diakinesis and metaphase I of both *him-8(e1489)* and *top-2(it7); him- 8(e1489)* (**Figure S1**). From these data we conclude that TOP-2 is specifically required for the maintenance of REC-8 cohesin but not COH-3/4.

### *top-2(it7)* mutants have disrupted AIR-2 recruitment

In late meiotic prophase I, the Aurora B kinase, AIR-2, phosphorylates REC-8 marking it for removal from the bivalent short arms (Ferrandiz et al., 2018; Kaitna et al., 2002; Rogers et al., 2002; Severson & Meyer, 2014). In meiosis II, AIR-2 marks REC-8 for removal from the sister chromatids. This two-step mechanism of cohesin removal is accomplished through the spatial and temporal restriction of AIR-2, which is first found on the bivalent short arms during meiosis I and between sister chromatids in meiosis II (Rogers et al., 2002). Because REC-8 appears to be prematurely removed from chromosomes in *top-2(it7)* spermatogenic germlines and AIR-2 is responsible for marking REC-8 for removal, we investigated whether the localization of AIR-2 is disrupted in these germlines. As previously described (Shakes et al., 2009), AIR-2 localized to the external surface of karyosomes and then relocalized to the short arm of diakinesis bivalents in *him-8(e1489)* spermatogenesis (**Figure 2A**). AIR-2 remains associated with segregating homologs during meiosis I, relocalizes between sister chromatids at metaphase II, and remains associated with the segregating chromosomes at anaphase II (**Figure 2A**). In *top-2(it7); him-8(e1489)* spermatogenesis AIR-2 was restricted to discrete areas of the chromosomes in the karyosome through metaphase I similar to controls. However, as chromosomes entered anaphase I, AIR-2 was ectopically localized with AIR-2 staining covering all regions of the chromosomes (global chromatin localization, **Figure 2A & C**). As the homologous chromosomes are unable to segregate in *top-2(it7); him-8(e1489)*, the chromosomes remain in an anaphase I configuration until the residual body is cleared (Jaramillo-Lambert et al., 2016). AIR-2 eventually disappears from the *top-2(it7); him-8(e1489)* anaphase I chromosomes (late anaphase I, **Figure 2A & C**).

**Figure 2.**
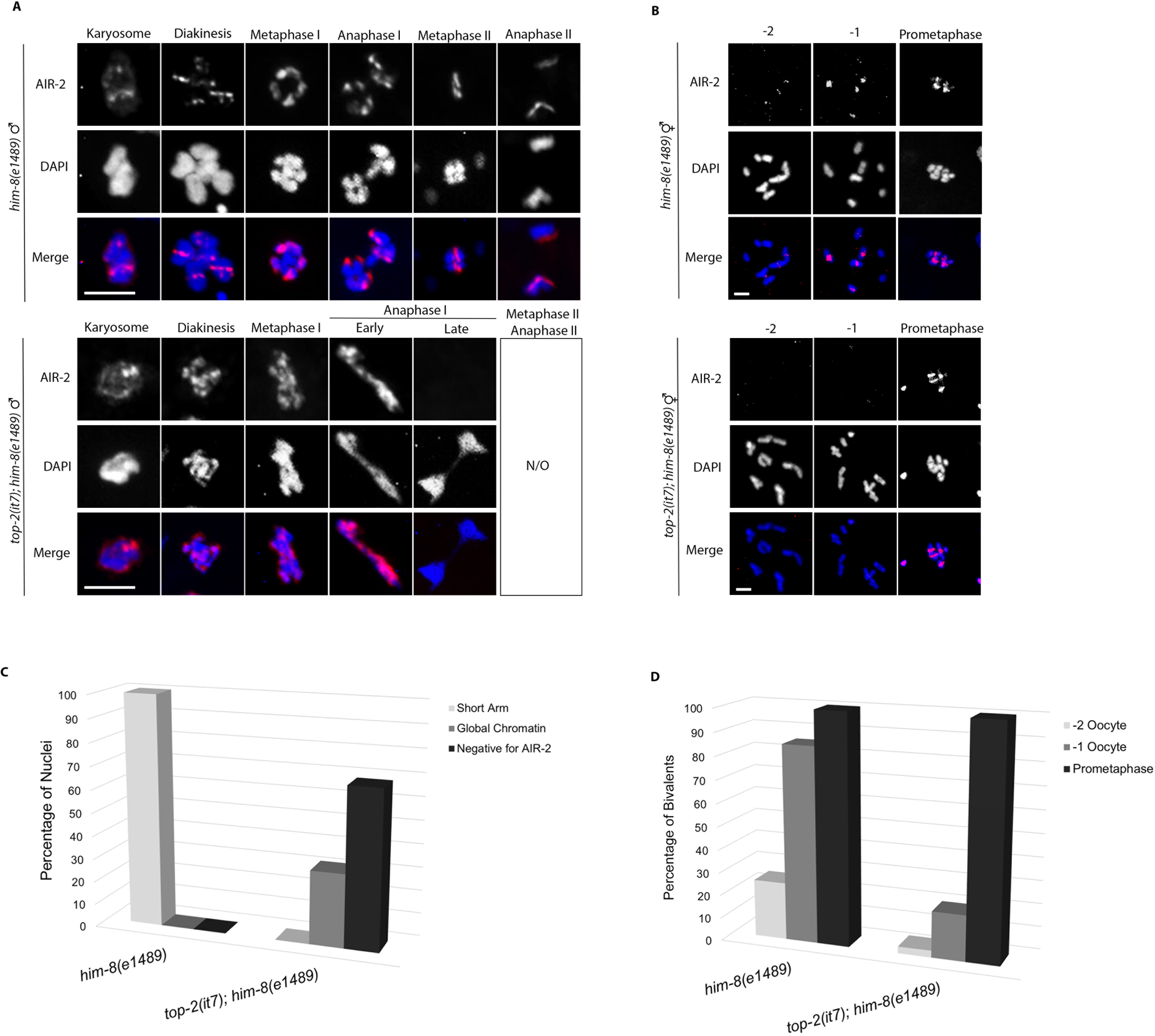
TOP-2 is required for proper localization of AIR-2 during both oogenesis and spermatogenesis. Immunostaining of AIR-2 (red) in control [*him-8(e1489)*] and *top-2(it7); him-8(e1489)* spermatogenesis (A) and oogenesis (B) counterstained with DAPI (blue). (A) Z-projections of karyosome through anaphase II nuclei in spermatogenesis. AIR-2 is localized to the DNA through anaphase II in control spermatogenesis. In *top-2(it7); him-8(e1489)* spermatogenesis AIR-2 is ectopically localized on anaphase I chromosomes. (B) Z-projections of diakinesis and prometaphase nuclei in oocytes. AIR-2 is localized to the short arm of diakinesis bivalents in the -1 oocyte and to prometaphase chromosomes in control animals. In *top- 2(it7); him-8(e1489),* AIR-2 is absent from the -1 oocyte, but localizes to chromosomes in prometaphase of meiosis I. (C) Quantification of the number of anaphase I nuclei either negative, short arm only, or ectopic [global chromatin] for AIR-2 localization during spermatogenesis in *him-8(e1489)* and *top-2(it7); him-8(e1489).* (D) Quantification of the number of diakinesis bivalents displaying AIR-2 localization in the - 2 and -1 oocytes and prometaphase I in *him-8(e1489)* and *top-2(it7); him-8(e1489)* oogenesis. N/O = not observed. Scale bar = 5 μm.

In diakinesis oocytes, AIR-2 localization is tightly regulated to restrict AIR-2 to the short arm of the bivalent of the two most proximal oocytes (-2 and -1 oocytes) and remains on the short arm through prometaphase I in the one-cell embryo. As we previously showed that the *top-2(it7)* mutant did not disrupt chromosome segregation during oogenesis, we hypothesized that AIR-2 would localize to the bivalent short arm in *top-2(it7)* oogenic germlines. Surprisingly, we found that in *top-2(it7)* AIR-2 fails to localize to the -2 and -1 oocytes, however it is present on prometaphase I chromosomes of the one-cell embryo (**Figure 2B & D**). These results demonstrate that the loss of TOP-2 function disrupts at least one protein in the SCC pathway of oogenesis. However, the loss of TOP-2 function effects proteins required for homologous chromosome segregation differently in spermatogenesis and oogenesis.

### TOP-2 is required for localization of sister chromatid cohesion release pathway proteins to prevent the untimely removal of REC-8 during meiosis I

Because the premature loss of REC-8 and ectopic localization of AIR-2 in *top-2(it7)* mutant spermatogenesis is reminiscent of defects in the SCC release pathway, we next examined the localization patterns of key SCC components. In oogenesis, REC-8 cohesin removal is both spatially and temporally regulated by the sister chromatid cohesion (SCC) release pathway (de Carvalho et al., 2008; Ferrandiz et al., 2018). To determine if this was also true in spermatogenesis, we examined the localization patterns of the proteins involved in SCC release in the male germline.

In diakinesis oocytes Haspin-mediated phosphorylation of Histone H3 on threonine 3 (H3pT3) cues the recruitment of AIR-2 to the bivalent short arm (Ferrandiz et al., 2018). We next sought to determine if the ectopic localization of AIR-2 in anaphase I of *top-2(it7)* spermatogenic germlines was due to ectopic phosphorylation of Histone H3 threonine 3. During spermatogenesis H3pT3 was detected at low levels starting at the karyosome stage with levels increasing through metaphase I (**Figure 3A & B**, **Figure S2A**). To help clarify the positional localization of H3pT3 on the diakinesis bivalents during spermatogenesis we double stained the germlines with the chromosomal axis component HTP-3, which localizes to the meiotic chromosome axes and is found on both the long and short arms of diakinesis bivalents in oocytes (Goodyer et al., 2008; Severson et al., 2009). In control germlines [*him-8(e1489)*] HTP-3 localizes to chromosome axes of pachytene chromosomes through the early karyosome (**Figure 3A & S2A**). HTP-3 then disassociates from the chromosome axes as they progress through the condensation zone, until it reassociates on the long and short arms of the bivalents at diakinesis and metaphase I (**Figure 3A & Figure S2A, S2C**). The cross-shaped localization pattern of HTP-3 at diakinesis can be more clearly observed in the single-plane images (**Figure S2D**). HTP-3 localization was no longer observed after metaphase I (**Figure 3A**). Double staining of HTP-3 and H3pT3 showed that H3pT3 localizes to the short arm of the bivalent at diakinesis through metaphase I (**Figure 3A**). H3pT3 localization is also found associated with the chromosomes in anaphase I, between sister chromatids in metaphase II, and with segregating sister chromosomes at anaphase II (**Figure 3A**).

**Figure 3.**
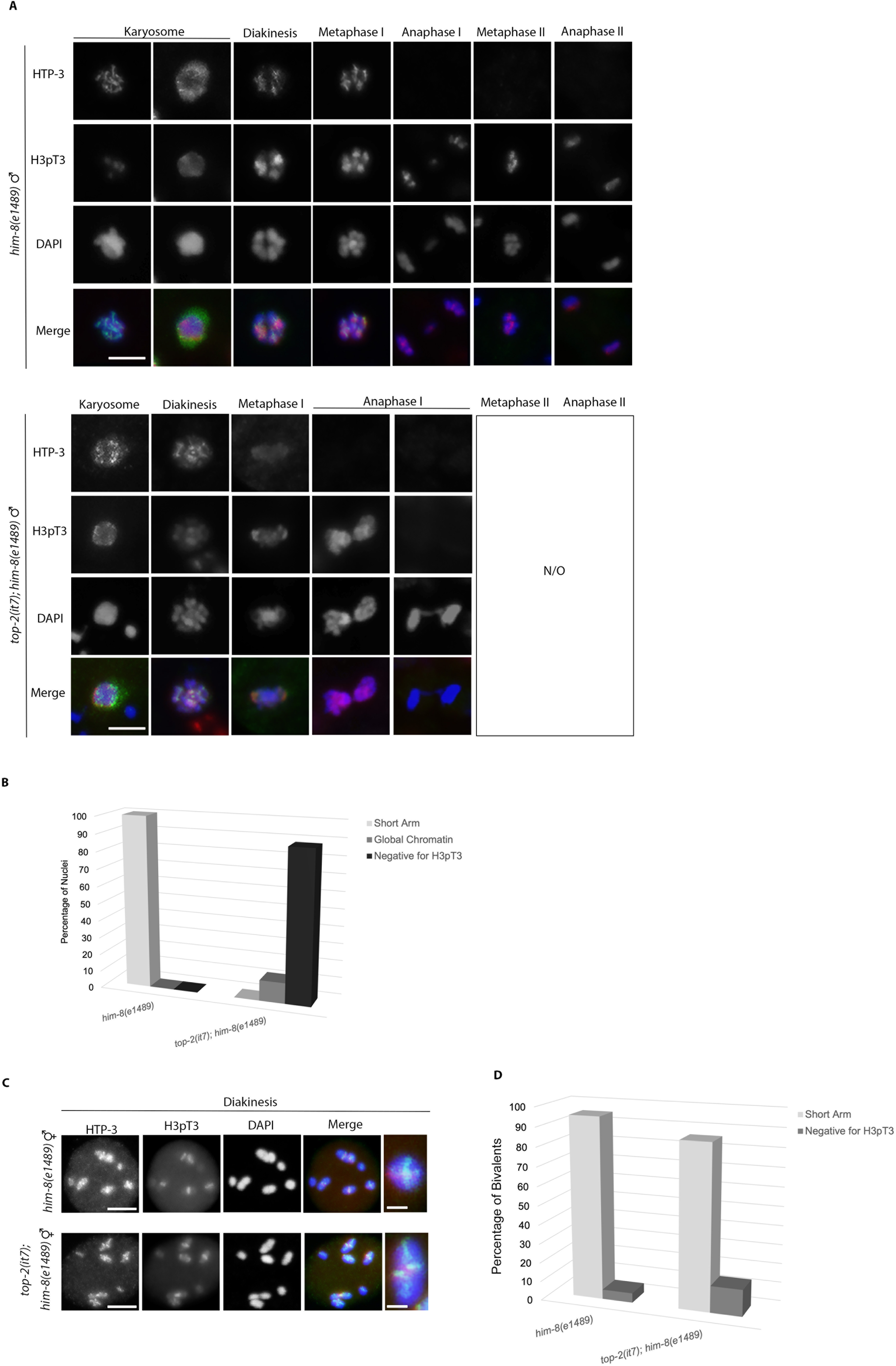
TOP-2 is required for the proper localization of H3 T3 phosphorylation and HTP-3 during spermatogenesis. Immunostaining of H3pT3 (red) and HTP-3 (green) in control [*him-8(e1489)*] and *top-2(it7); him-8(e1489)* in spermatogenesis (A) and oogenesis (C), counterstained with DAPI (blue). A) Z-projections of karyosome through anaphase II nuclei in spermatogenesis. H3pT3 localizes to the DNA at the karyosome where it remains through anaphase II in *him-8(e1489)* spermatogenesis. In *top-2(it7); him-8(e1489)* spermatogenesis H3pT3 loads properly at the karyosome but is ectopically localized at anaphase I. HTP-3 is localized to the chromosome axes at the karyosome, disassociates in the condensation zone, and reassociates with chromosomes at diakinesis where it remains through metaphase I in control germlines. While HTP-3 localization was detected in *top-2(it7); him-8(e1489)* spermatogenesis in the karyosome stage through metaphase I, the localization pattern was perturbed. (B) Quantification of the number of anaphase nuclei either negative, short arm only, or ectopic [global chromatin] for H3pT3 localization during control and *top-2(it7); him-8 (e1489)* spermatogenesis. (C) Z-projections of diakinesis nuclei in the -1 oocyte, and a single magnified bivalent in the last column. H3pT3 is localized to the short arm of diakinesis bivalents in the -1 oocyte in control and *top-2(it7); him-8(e1489)* animals. HTP-3 is localized to both the long and short arms of diakinesis bivalents in the -1 oocyte in control and *top-2(it7); him-8(e1489)* animals. (D) Quantification of the number of diakinesis bivalents with H3pT3 localization on the short arm only, or negative for H3pT3 in either negative or short arm only. N/O = not observed. Scale bar = 5 μm. Magnified bivalent scale bar = 2 μm.

In *top-2(it7); him-8(e1489)* spermatogenic germlines HTP-3 localizes to chromosome axes in pachytene (**Figure S2B**) but displays a disrupted chromosomal association at the karyosome and in diakinesis nuclei (**Figure 3A**). Cross-shaped HTP- 3 structures are observed in diakinesis nuclei, but these do not appear to be associated with the chromosomes (**Figure S2D**). H3pT3 was found associated with chromosomes in the karyosome and diakinesis nuclei with some brighter H3pT3 foci at diakinesis. However, in metaphase I and anaphase I nuclei H3pT3 localization was disrupted. H3pT3 had an ectopic localization pattern on early anaphase structures and was absent from late anaphase I structures, similar to AIR-2 localization in *top-2(it7); him-8(e1489)* spermatogenic germlines (**Figure 3A & 3B**). From this we conclude that ectopic H3pT3 localization leads to ectopic AIR-2 localization and thus the precocious removal of REC- 8 in *top-2(it7)* mutant spermatogenesis.

We also examined the localization of HTP-3 and H3pT3 in *top-2(it7); him- 8(e1489)* diakinesis oocytes. Neither HTP-3 nor H3pT3 localization was disrupted in the mutant germlines. HTP-3 localized to both the long and short arm of the bivalent and H3pT3 localized to the short arm (**Figure 3C & D, Figure S2E**). Thus, *top-2(it7)* is not affecting timing of AIR-2 recruitment in oocytes at the level of H3 phosphorylation.

The stepwise removal of cohesin in oogenesis is mediated through the spatiotemporal regulation of AIR-2, which in turn is regulated through the spatial recruitment of several proteins starting with the axis components HTP-1 and HTP-2 [82% identical and referred to as HTP-1/2 (Martinez-Perez et al., 2008)]. The N-terminal domain of HTP-1/2 acts as a scaffold through LAB-1 to promote the recruitment of protein phosphatase 1 (PP1) to selectively dephosphorylate H3pT3 on the bivalent long arm (Ferrandiz et al., 2018). We next asked whether localization of the scaffold proteins HTP-1/2 were disrupted in *top-2(it7)* mutant germlines. In control [*him-8(e1489)*] animals, we found that HTP-1/2 localized to the chromosomes in the transition zone (leptotene and zygotene) through pachytene and diakinesis in prophase I and then was lost from metaphase I chromosomes during spermatogenesis (**Figure 4A, B & Figure S3**). In *top-2(it7); him-8(e1489)* spermatogenic germlines we found that HTP-1/2 localized to chromosomes similarly to control germlines in early stages of meiotic prophase I (**Figure 4A, B & Figure S3**). However, whereas HTP-1/2 was detected on 100% of diakinesis nuclei in control germlines, 50% of diakinesis nuclei lacked HTP-1/2 in *top-2(it7); him-8(e1489)* germlines (**Figure 4A & B**). Examination of HTP-1/2 localization in both control [*him-8(e1489)*] and *top-2(it7); him-8(e1489)* oogenic germlines found that HTP-1/2 localizes to the bivalent long arm (**Figure 4C & D**). These data indicate that TOP-2 is required to maintain HTP-1/2 localization during spermatogenesis, but not during oogenesis.

**Figure 4.**
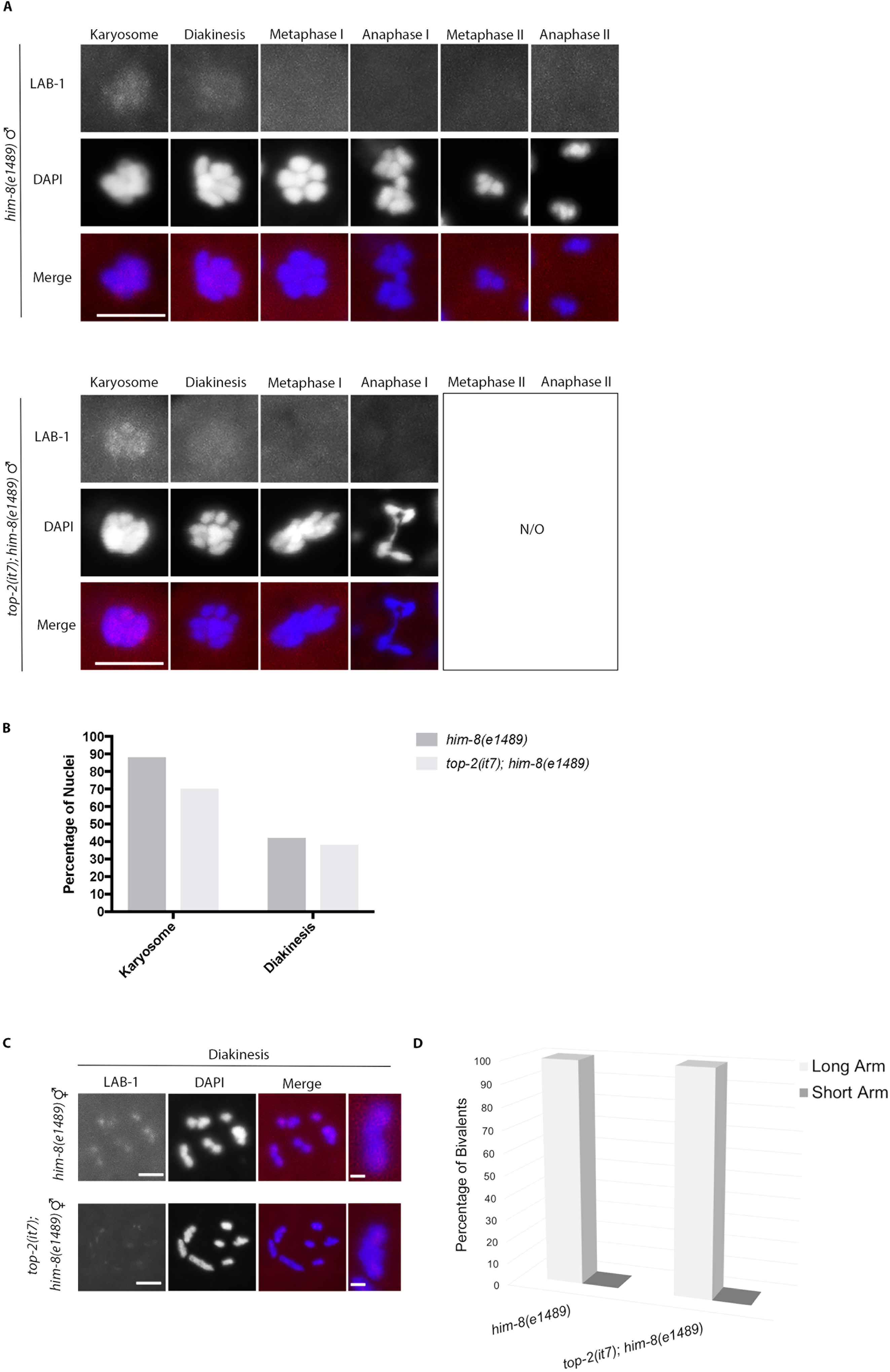
TOP-2 is required for proper localization of HTP-1/2 during spermatogenesis, but not oogenesis. Immunostaining of HTP-1/2 (red) in control [*him-8(e1489)*] and *top-2(it7); him-8(e1489)* in spermatogenesis (A) and oogenesis (C), counterstained with DAPI (blue). (A) Z-projections of karyosome through anaphase II nuclei in spermatogenesis. In control animals, HTP-1/2 is localized to the DNA through diakinesis stage nuclei and is removed at metaphase I. In *top-2(it7); him-8(e1489)* spermatogenesis, HTP-1/2 shows proper localization through the karyosome stage, however it is prematurely removed at diakinesis. (B) Quantification of the percent nuclei positive for HTP-1/2 localization in the karyosome, diakinesis, and metaphase I of meiosis I during spermatogenesis. (C) HTP-1/2 localization at diakinesis in the -1 oocyte of *him-8(e1489)* and *top-2(it7); him-8(e1489)* oogenesis with a single magnified bivalent in the last column. (D) Quantification of the percent bivalents with HTP-1/2 localization on the long arm or the short arm of diakinesis oocytes in both *him-8(e1489)* and *top- 2(it7); him-8(e1489)*. N/O = not observed. Scale bar = 5 μm. Magnified bivalent scale bar = 2 μm.

We also examined another component of the oocyte SCC release pathway in spermatogenesis, LAB-1. LAB-1 interacts with PP1 to restrict H3pT3 to the short arms of late diakinesis oocyte bivalents (de Carvalho et al., 2008; Ferrandiz et al., 2018). As LAB-1, which localizes to the long arms of oocyte bivalents, serves as the docking site for PP1 phosphatase, we asked if the ectopic localization of H3pT3 in spermatogenesis was due to either a failure to localize LAB-1 to chromosomes or mislocalization of the protein. In control spermatogenic germlines LAB-1 was detected from late pachytene through the condensation zone (**Figure 5A & Figure S4**). LAB-1 associates with chromosomes during late pachytene of meiotic prophase I (**Figure S4A & S4B**). LAB-1 chromosome tracks are detected through diplotene and then dissociate from the chromosomes throughout the condensation zone (diplotene through karyosome stages) until it is no longer detected at diakinesis (**Figure 5A & S4B**). This is distinct from oogenesis where LAB-1 is first detected in germ cells entering meiotic prophase I (leptotene/zygotene) until late meiotic prophase I (diakinesis) (de Carvalho et al., 2008). In *top-2(it7); him-8(e1489)* mutant spermatogenesis LAB-1 localizes to chromosomes similarly to the localization pattern observed in control germlines with no major disruption to localization detected (**Figure 5A & B**). Examination of the localization of LAB-1 in *top-2(it7); him-8(e1489)* diakinesis oocytes found that LAB-1 localized to the long arm of the bivalents in the mutant germlines similar to control germlines (**Figure 5C & D**).

**Figure 5.**
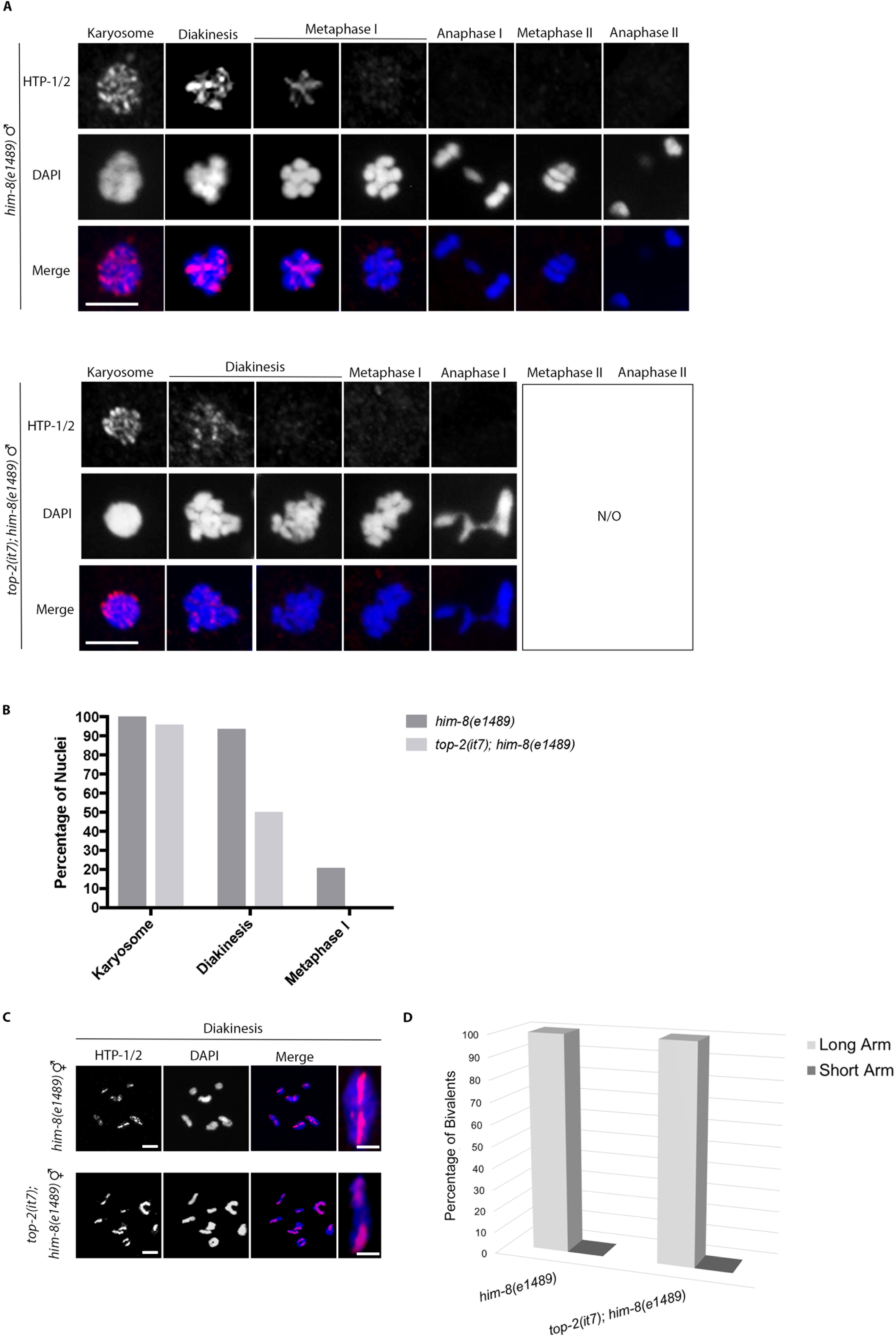
TOP-2 is not required for proper localization of LAB-1 during spermatogenesis and oogenesis. Immunostaining of LAB-1 (red) in control [*him- 8(e1489)*] and *top-2(it7); him-8(e1489)* in spermatogenesis (A) and oogenesis (C), counterstained with DAPI (blue). (A) Z-projections of karyosome through anaphase II nuclei in spermatogenesis. In control animals LAB-1 is localized to the DNA at the karyosome and is removed at diakinesis stage nuclei in both control and *top-2(it7); him- 8(e1489)* animals. (B) Quantification of the percent nuclei positive for LAB-1 at the karyosome and diakinesis stages of meiosis I during spermatogenesis. (C) Z-projections of LAB-1 localization at diakinesis in the -1 oocyte of control and *top-2(it7); him-8(e1489)* animals. A single magnified bivalent is presented in the last column. LAB- 1 localizes to the long arm in both control and *top-2(it7); him-8(e1489)* diakinesis chromosome bivalents. (D) Quantification of the percentage of diakinesis chromosome bivalents with LAB-1 localization on the long arm or short arm. N/O = not observed. Scale bar = 5 μm. Magnified bivalent scale bar = 2 μm.

As LAB-1 appears to have a sex-specific pattern of localization with LAB-1 appearing much later during spermatogenesis, it is possible that LAB-1 does not play a major role in the SCC pathway during spermatogenesis. To address this possibility, we counted the number of DAPI staining bodies in diakinesis nuclei in *lab-1(tm1791)* mutant spermatogenic germlines. *C. elegans* males have five pairs of autosomes and one sex chromosome (the X chromosome), which are observed as six DAPI staining bodies in wild type. Analysis of *lab-1(tm1791)* worms revealed the presence of univalents in diakinesis spermatocytes (up to 10 DAPI staining bodies) (**Figure SF4**), indicating premature separation of homologous chromosomes. However, mating *lab- 1(tm1791)* males with *fog-2(oz40)* females did not result in a significant decrease in progeny viability [N2: 98.0%, *lab-1(tm1791)*: 91.8%, p=0.07] suggesting that although unpaired, the univalents are distributed correctly to make viable haploid gametes. These data demonstrate that LAB-1 is involved in the proper pairing of homologous chromosomes during spermatogenesis. Nevertheless our LAB-1 immunostaining results suggest that in *top-2(it7)* spermatogenic germlines, it is not a mislocalization of LAB-1 that is causing precocious cohesin removal. Additional studies are needed to fully determine the role of LAB-1 in spermatogenesis.

### Reduction of *wee-1.3* restores AIR-2 recruitment to diakinetic chromosomes

During oogenesis AIR-2 is not only regulated through restricted spatial recruitment to chromosomes, AIR-2 localization is also temporally regulated. CDK-1 and cyclin B, as part of the maturation promoting factor (MPF), control the switch between diakinesis and metaphase I (Doree & Hunt, 2002). In *C. elegans* oogenesis, CDK- 1/MPF controls the timing of AIR-2 recruitment to the -2 and -1 oocytes (Ferrandiz et al., 2018). Wee1 kinase mediates the inhibition of Cdk1 through the phosphorylation of specific inhibitory sites (Doree & Hunt, 2002). Thus, WEE-1.3 regulates the timing of AIR-2 recruitment to meiotic bivalents through the inhibition of MPF (Ferrandiz et al., 2018). As we observed a lack of AIR-2 staining in the -2 and -1 oocytes of *top-2(it7); him-8(e1489)* oogenic germlines, we investigated if the timing of AIR-2 localization could be restored in mutant germlines through inhibition of the WEE-1.3 kinase. RNAi of *wee-1.3* restored AIR-2 localization to the bivalent short arm of *top-2(it7); him-8(e1489)* diakinesis chromosomes of the -1 oocytes (**Figure 6A & B**). This suggests that TOP-2 may help regulate the timing of AIR-2 recruitment during *C. elegans* oogenesis.

**Figure 6.**
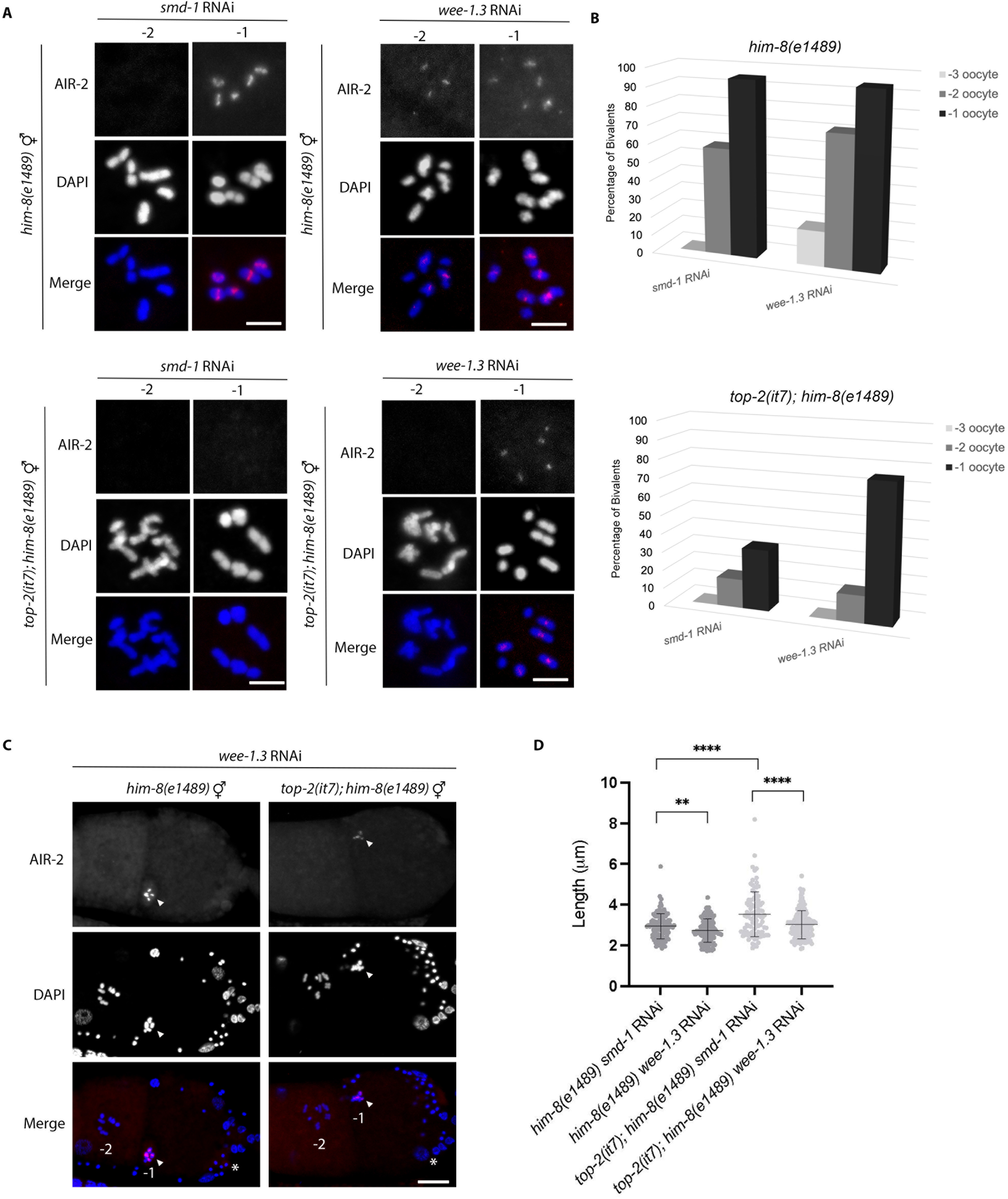
Reduction of WEE-1.3 restores AIR-2 recruitment to diakinetic chromosomes during oogenesis. Immunostaining of AIR-2 (red) and DAPI (blue) in control [*him-8(e1489)*] and *top-2(it7); him-8(e1489)* treated with control and *wee-1.3* RNAi during oogenesis. (A) Z-projections of -1 and -2 diakinesis oocytes, post treatment with control RNAi [*smd-1,* left panels] or *wee-1.3* RNAi [right panels]. In control animals treated with *smd-1* RNAi, AIR-2 localizes to the short arm in the most proximal oocyte. Control worms treated with *wee-1.3* RNAi have precocious recruitment of AIR-2 in the - 2 oocyte. *top-2(it7); him-8(e1489)* animals treated with *smd-1* RNAi fail to load AIR-2 to the short arm of diakinesis chromosome bivalents. *top-2(it7); him-8(e1489)* treated with *wee-1.3* RNAi restores AIR-2 localization to the -1 oocyte. (B) Quantification of the percent of bivalents with AIR-2 localization on the short arm of the -1, -2, and –3 oocytes in control and *top-2(it7); him-8(e1489)* animals treated with *smd-1* or *wee-1.3* RNAi. (C) Z-projection of the -1 and -2 oocytes in control and *top-2(it7); him-8(e1489)* worms treated with *wee-1.3* RNAi. Arrow points to the more condensed chromosome structure observed in *wee-1.3* depleted oocytes, and asterisks mark the spermatheca. (D) Quantification of -1 oocyte chromosome length measurements in *him-8(e1489)* and *top-2(it7); him-8(e1489)* hermaphrodites treated with control (*smd-1*) or *wee-1.3* RNAi. Chromosome length is significantly decreased in *top-2(it7); him-8(e1489)* worms treated with *wee-1.3* RNAi when compared to control RNAi. Statistical analysis performed using a t-test. ****p = <0.0001, **p = <0.01. Scale bar = 5 μm.

TOP-2 has been shown to be involved in chromosome condensation and in setting chromosome length in the soma (Ladouceur et al., 2017; Nitiss, 2009). We reasoned that the *top-2(it7)* mediated disruptions to the localization of SCC pathway proteins could be due to disruptions in chromosome structure (condensation, length). Indeed, during the immunostaining experiments it appeared that the diakinesis chromosomes of *top-2(it7); him-8(e1489)* oocytes were longer than *him-8(e1489)* control oocytes. To determine if *top-2(it7)* affects chromosome length, we measured the lengths of individual bivalents in *him-8(e1489)* vs. *top-2(it7); him-8(e1489)* treated with control RNAi (*smd-1*). We found that chromosome length was increased in *top-1(it7); him-8(e1489); smd-1* RNAi compared to *him-8(e1489); smd-1* RNAi control (3.53 μm vs2.95 μm, p<0.0001, **Figure 6D**). Depletion of *wee-1.3* via RNAi results in hypercompaction of oocyte chromosomes [*him-8(e1489) smd-1* RNAi: 2.95 μm vs *him- 8(e1489) wee-1.3* RNAi: 2.74 μm, p<0.01, **Figure 6C-D**, (Allen et al., 2014; Burrows et al., 2006)]. This suggests that the rescue of AIR-2 localization in *top-2(it7); him- 8(e1489)* oocytes depleted of *wee-1.3* could be due to changes in chromosome structure caused by the premature activation of the cell cycle. We measured chromosome length of oocyte diakinesis bivalents in *top-2(it7); him-8(e1489)* treated with *smd-1* RNAi vs. *wee-1.3* RNAi. Chromosome length was decreased in *top-2(it7); him-8(e1489) wee-1.3* RNAi (3.03 μm) vs. *top-2(i7); him-8(e14890 smd-1* RNAi (3.53 μm) (p<0.0001, **Figure 6D**). From these experiments we conclude that TOP-2 is required for proper chromosome structure/length.

## DISCUSSION

The accurate segregation of homologous chromosomes during meiotic prophase I is a complex process. In meiotic prophase I, homologous chromosomes pair, synapse, and recombine. These intricate interactions require proteins that load cohesins and establish SCC, maintain cohesin binding, and mediate the regulated two-step removal of SCC from chromosomes. Previous studies have elegantly determined the mechanisms that control the establishment and release of SCC in *C. elegans* oogenesis (Ferrandiz et al., 2018; Rogers et al., 2002; Severson et al., 2009; Severson & Meyer, 2014; Tzur et al., 2012). Our observation that a *top-2* hypomorphic mutation causes precocious loss of REC-8 from chromosomes during spermatogenesis prompted this investigation into SCC release in the male germline. We found that the SCC release pathway components in *C. elegans* spermatogenesis have similar localization patterns to those in *C. elegans* oogenesis. At diakinesis first the HORMA-domain proteins HTP-1/2, which localize along chromosome axes in pachytene, are redistributed to localize to the bivalent long arm. We also found that the meiosis-specific cohesin REC-8 localizes to the long arm of diakinesis bivalents. In contrast, COH-3/4 (additional meiosis-specific cohesins), AIR-2, and H3 T3 phosphorylation are all found on the short arm of diakinesis bivalents. Interestingly, we found that LAB-1, the PP1 phosphatase binding partner, localizes to chromosomes in pachytene through the karyosome stages, but is largely absent from diakinesis chromosomes. This differs from SCC release in *C. elegans* oocytes, which require LAB-1 for PP1 recruitment to the bivalent long arm at diakinesis. PP1 antagonizes H3 T3 phosphorylation on the long arm thus restricting H3 T3 phosphorylation and AIR-2 localization to the bivalent short arm. These findings reveal that while localization of components required for SCC release pathway are similar in spermatogenesis and oogenesis, the localization of these components in the male and female germline are not identical.

In addition to differences in SCC release between oogenesis and spermatogenesis, this study revealed that the enzyme Topoisomerase II is differentially required for SCC release in the germlines of males and females. The male germline is much more sensitive to a reduction in Topoisomerase II function. When males harboring a temperature-sensitive allele of *topoisomerase II*, *top-2(it7)*, were incubated at the nonpermissive temperature, several SCC components were disrupted. HTP-1/2 were prematurely removed from the long arms of diakinesis bivalents, leading to ectopic localization of H3 T3 phosphorylation, ectopic AIR-2 localization, and the precocious removal of REC-8 cohesin. In contrast, *top-2(it7)* oogenic germlines displayed normal localization of most SCC components. The exception was AIR-2, which failed to localize to diakinesis oocytes, but was present on prometaphase I bivalent short arms in the one cell embryo.

As homologous chromosomes prepare to segregate at anaphase of meiosis I, the chromosomes become highly condensed and restructured during late meiotic prophase to form the cruciform shape of the bivalent. Chromosome condensation and remodeling happen in both spermatogenesis and oogenesis, therefore how does oogenesis compensate for the disrupted localization of AIR-2 in the *top-2* mutant and allow for accurate segregation of homologous chromosomes at meiosis I? One major difference between late meiotic prophase in oogenesis and spermatogenesis is timing of these events. In oogenesis the transition from diakinesis to metaphase I is temporally regulated by the MPF. The MPF activates many hallmark events including nuclear envelope breakdown, rearrangement of the cortical cytoskeleton, and chromosome condensation and congression (Jones, 2004; Von Stetina & Orr-Weaver, 2011). In *top- 2(it7)* mutants, we observed a lack of AIR-2 staining on -1 oocytes (diakinesis), but, surprisingly, AIR-2 was robustly localized to prometaphase I chromosomes (**Figure 2**). At this stage of the meiotic cell cycle the chromosomes are in their most condensed configuration. Thus, we hypothesize that chromosome condensation state might also regulate the timing of AIR-2 localization (**Figure 7**). Indeed, we found that knock-down of the MPF inhibitor kinase WEE-1.3, which results in precocious oocyte maturation including chromosome hypercondensation and congression (Allen et al., 2014; Burrows et al., 2006), restores AIR-2 localization to the -1 oocytes of *top-2(it7)* (**Figure 6**). Our chromosome length data also support this hypothesis. Oocyte diakinesis chromosome lengths are significantly increased in *top-2(it7); him-8(e1489)*; *smd-1* RNAi compared to *him-8(e1489); smd-1* RNAi and AIR-2 localization is restored on the shorter chromosomes of *top-2(it7); him-8(e1489); wee-1.3* RNAi (**Figure 6C**). From these data we propose that TOP-2 plays a role in chromosome structure in oogenic germlines, but relatively small disruptions caused by *top-2(it7)* can be compensated by later events of the diakinesis to metaphase transition (**Figure 7**).

**Figure 7.**
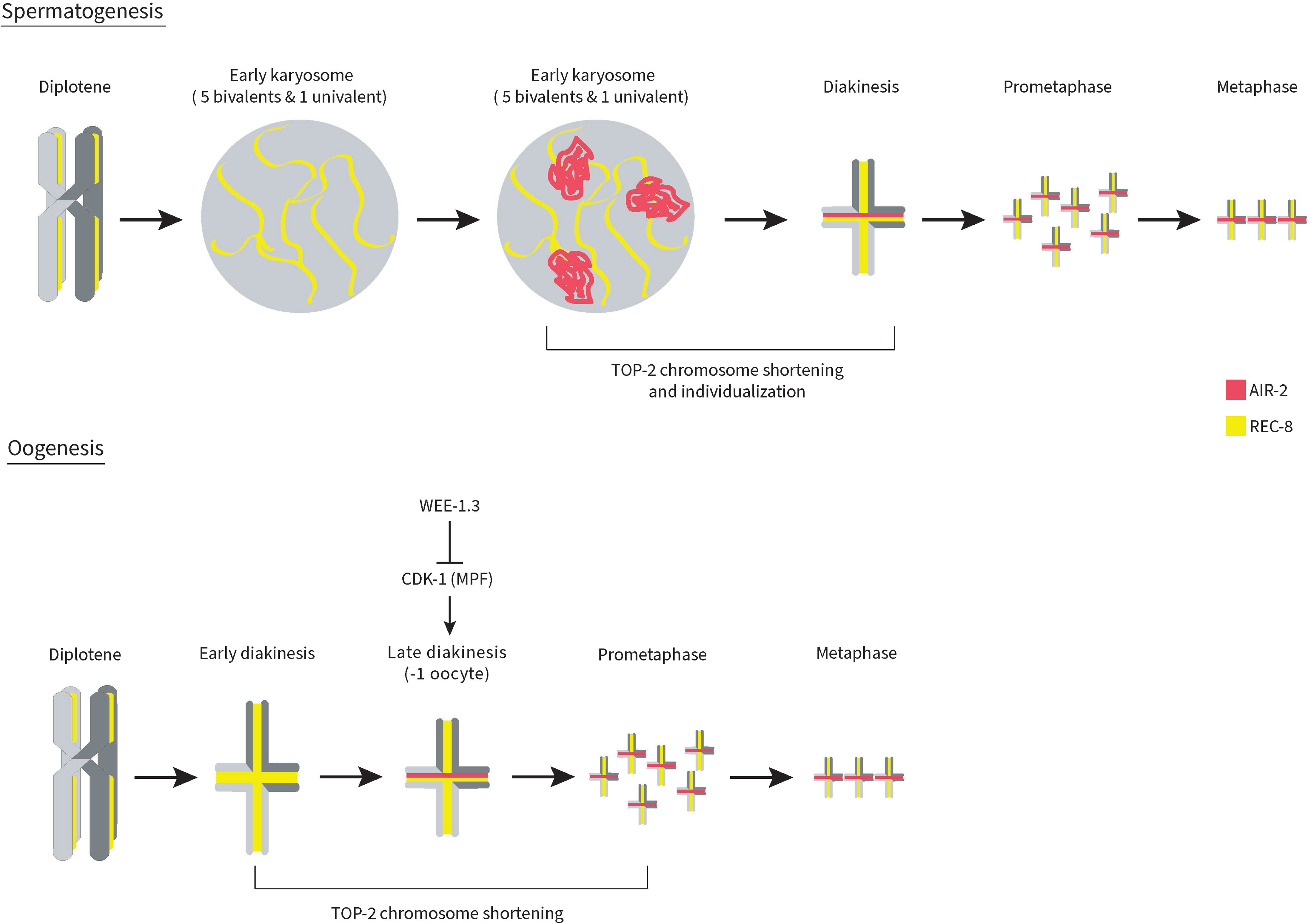
Model of TOP-2 function in the events that control the release of SCC and chromosome remodeling in spermatogenesis and oogenesis. During spermatogenesis, TOP-2-mediated chromosome remodeling is required at the late karyosome to diakinesis transition to individualize and compact (shortening and condensation) the chromosomes. This allows for the proper maintenance of chromosome axis components (e.g. HTP-1/2, not depicted) and the subsequent regulation of the spatial localization of AIR-2 to the short arm of diakinesis bivalents and the timely release of REC-8. During oogenesis, TOP-2 functions at diakinesis through prometaphase I to mediate chromosome compaction (shortening and condensation). Once the chromosomes have reached the correct amount of compaction and in conjunction with the MPF, AIR-2 can be recruited to the short arms of the bivalents.

### What is the link between TOP-2, chromosome structure, and the maintenance of HTP-1/2 localization in spermatogenesis?

In late meiotic prophase, chromosomes of both oocytes and spermatocytes resolve and condense as they prepare for the meiotic divisions. However, unlike oocytes which proceed from diakinesis to metaphase only after activation by MPF, spermatocytes proceed directly from diakinesis through the meiotic divisions (Chu & Shakes, 2013). Another difference in meiotic prophase between oogenesis and spermatogenesis, is that as spermatogenic chromosomes start to undergo chromatin remodeling in late prophase, the chromosomes condense and coalesce into a single mass called a karyosome (Shakes et al., 2009). It has been previously proposed that homologous chromosome compaction involves the cooperative actions of cohesins, condensins, and DNA topoisomerases (Cahoon & Libuda, 2019; Hillers et al., 2017; Kleckner, 2006; Uhlmann, 2016). Our results support this hypothesis in spermatogenesis; loss of TOP-2 function in spermatogenesis leads to the premature loss of the meiotic cohesin REC-8 (**Figure 1**). We found that the loss of REC-8 is caused by the upstream loss of the chromosome axis components HTP-1/2. How does loss of *top-2* function cause HTP-1/2 to be precociously lost from meiotic chromosomes? We put forward the model that TOP-2-induced chromatin remodeling is required to prepare the chromosomes into a shape that promotes the continued binding of specific chromosome axis components (**Figure 7**). Several examples in multiple organisms link Topoisomerase II and chromosome structure (Hartsuiker et al., 1998; Hughes & Hawley, 2014; Li et al., 2013; Liang et al., 2015; Maeshima & Laemmli, 2003; Mengoli et al., 2014; Xu & Manley, 2007; Zhang et al., 2014). In addition, a connection between Topoisomerase II and meiotic axis components was demonstrated in budding yeast (Heldrich et al., 2020). In yeast, temperature-sensitive *top-2* mutants have delayed removal of the chromosome axis protein Hop1 (Heldrich et al., 2020). HTP-1/2 are homologs of yeast Hop1 (Martinez-Perez & Villeneuve, 2005). In contrast to budding yeast, we found that a temperature-sensitive *top-2* mutant leads to the early removal of HTP-1/2 from chromosomes in *C. elegans* spermatogenesis (**Figure 4**). This is likely due to different mechanisms of homolog remodeling in yeast and worms. In budding yeast, Hop1 removal prior to SC assembly is required to recruit Pch2/TRIP13 for the remodeling of synapsed homologous chromosomes (Heldrich et al., 2020). In contrast, *C. elegans* HTP-1/2 are required to remain on chromosomes much later in meiotic prophase to promote the accurate localization of SCC release proteins. HTP-1/2 recruitment to chromosomes in early prophase is dependent on either binding axis proteins HTP-3 or HIM-3. HIM-3 is also dependent on HTP-3 for recruitment to chromosome axes (Kim et al., 2014). Although we found that HTP-3 localizes to late prophase nuclei in *top-2(it7)* spermatogenesis, HTP-3 appears to form aberrant structures in diakinesis/metaphase and in some cases does not even appear to be associated with chromatin (**Figure S2**).

One possibility is that TOP-2 directly binds to one or several of the axis proteins and loss of TOP-2 localization to chromosomes as in the *top-2(it7)* mutant (Jaramillo-Lambert et al., 2016) disrupts localization of other chromosome axis components. As disruption of HTP-3, HTP-1/2, and REC-8 localization were not observed in *top-2(it7)* oogenesis we do not believe this is likely. Another possibility is that TOP-2 is required to help restructure the chromosomes as they emerge from the aggregated chromatin mass of the karysome to become individualized entities of diakinesis nuclei. We believe this is the most plausible scenario. In *top-2(it7)* mutant spermatogenesis, chromosome structure appears grossly normal prior to the karyosome stage, but as the nuclei progress into diakinesis chromosome structure is severely disrupted in *top-2(it7)* mutant germlines with individual chromosome pairs indistinguishable from each other. Diakinesis chromosomes in oogenesis are also affected by the *top-2(it7)* mutant; bivalents are longer compared to controls. The more severe phenotype found in spermatogenesis (SCC disruption and chromosome segregation defects) is probably due to an increased requirement for TOP-2 in the karyosome to diakinesis transition (**Figure 7**).

In this study we focused on the role of topoisomerase II and SCC release in late meiotic prophase. While we cannot discount a role for topoisomerase II in early meiotic events, we did not observe any obvious defects prior to late meiotic prophase with the *top-2(it7)* mutant [This study and (Jaramillo-Lambert et al., 2016)]. In the future, identification of new alleles or utilization of a degron-tagged TOP-2 protein may reveal roles for TOP-2 in early meiotic events as well. New alleles will also be useful in answering how loss of TOP-2 affects chromosome remodeling in late meiotic prophase in both spermatogenesis and oogenesis.

## ACKNOWLEDGMENTS

We thank the following people for their gifts of antibodies: Jill Schumacher (AIR-2), Aaron Severson (COH-3/4), Monica Colaiacovo (LAB-1), and Abby Dernburg (HTP-1/2 and HTP-3). We thank Enrique Martinez-Perez for sharing strains. We also thank Amber Krauchunas and members of the Jaramillo-Lambert laboratory for critical reading of this manuscript. Microscopy access was supported by grants from the NIH-NIGMS (P20 GM103446), the NSF (IIA-1301765) and the State of Delaware. The LSM880 confocal microscope was acquired with a shared instrumentation grant S10 OD016361. Some strains were provided by the CGC, which is funded by NIH Office of Research Infrastructure Programs (P40 0D010440). This work was supported by National Institutes of Health R03HD098244 (A.J.L). C.R. was supported by NIH T32GM133395 and a University of Delaware Graduate Scholars Award.

## LITERATURE CITED

Allen, A. K., Nesmith, J. E., & Golden, A. (2014). An RNAi-based suppressor screen identifies interactors of the Myt1 ortholog of *Caenorhabditis elegans*. G3 (Bethesda, Md.), 4(12), 2329–2343. 10.1534/g3.114.013649 [doi]

Barber, T. D., McManus, K., Yuen, K. W., Reis, M., Parmigiani, G., Shen, D., Barrett, I., Nouhi, Y., Spencer, F., Markowitz, S., Velculescu, V. E., Kinzler, K. W., Vogelstein, B., Lengauer, C., & Hieter, P. (2008). Chromatid cohesion defects may underlie chromosome instability in human colorectal cancers. Proceedings of the National Academy of Sciences of the United States of America, 105(9), 3443–3448. 10.1073/pnas.0712384105 [doi]

Blat, Y., Protacio, R. U., Hunter, N., & Kleckner, N. (2002). Physical and functional interactions among basic chromosome organizational features govern early steps of meiotic chiasma formation. Cell, 111(6), 791–802. S0092867402011674 [pii]

Burrows, A. E., Sceurman, B. K., Kosinski, M. E., Richie, C. T., Sadler, P. L., Schumacher, J. M., & Golden, A. (2006). The *C. elegans* Myt1 ortholog is required for the proper timing of oocyte maturation. Development (Cambridge, England), 133(4), 697–709. dev.02241 [pii]

Cahoon, C. K., & Libuda, D. E. (2019). Leagues of their own: sexually dimorphic features of meiotic prophase I. Chromosoma, 128(3), 199–214. 10.1007/s00412-019-00692-x [doi]

Chu, D. S., & Shakes, D. C. (2013). Spermatogenesis. Advances in Experimental Medicine and Biology, 757, 171–203. 10.1007/978-1-4614-4015-4_7 [doi]

de Carvalho, C. E., Zaaijer, S., Smolikov, S., Gu, Y., Schumacher, J. M., & Colaiacovo, M. P. (2008). LAB-1 antagonizes the Aurora B kinase in *C. elegans*. Genes & Development, 22(20), 2869–2885. 10.1101/gad.1691208 [doi]

Doree, M., & Hunt, T. (2002). From Cdc2 to Cdk1: when did the cell cycle kinase join its cyclin partner? Journal of Cell Science, 115(Pt 12), 2461–2464.

Ferrandiz, N., Barroso, C., Telecan, O., Shao, N., Kim, H. M., Testori, S., Faull, P., Cutillas, P., Snijders, A. P., Colaiacovo, M. P., & Martinez-Perez, E. (2018). Spatiotemporal regulation of Aurora B recruitment ensures release of cohesion during *C. elegans* oocyte meiosis. Nature Communications, 9(1), 834–5. 10.1038/s41467-018-03229-5 [doi]

Gomez, R., Viera, A., Berenguer, I., Llano, E., Pendas, A. M., Barbero, J. L., Kikuchi, A., & Suja, J. A. (2014). Cohesin removal precedes topoisomerase II alpha-dependent decatenation at centromeres in male mammalian meiosis II. Chromosoma, 123(1- 2), 129–146. 10.1007/s00412-013-0434-9 [doi]

Goodyer, W., Kaitna, S., Couteau, F., Ward, J. D., Boulton, S. J., & Zetka, M. (2008). HTP-3 links DSB formation with homolog pairing and crossing over during *C. elegans* meiosis. Developmental Cell, 14(2), 263–274. 10.1016/j.devcel.2007.11.016 [doi]

Harper, N. C., Rillo, R., Jover-Gil, S., Assaf, Z. J., Bhalla, N., & Dernburg, A. F. (2011). Pairing centers recruit a Polo-like kinase to orchestrate meiotic chromosome dynamics in *C. elegans*. Developmental Cell, 21(5), 934–947. 10.1016/j.devcel.2011.09.001 [doi]

Hartsuiker, E., Bahler, J., & Kohli, J. (1998). The role of topoisomerase II in meiotic chromosome condensation and segregation in *Schizosaccharomyces pombe*. Molecular Biology of the Cell, 9(10), 2739–2750. 10.1091/mbc.9.10.2739 [doi]

Hassold, T., & Hunt, P. (2001). To err (meiotically) is human: the genesis of human aneuploidy. Nature Reviews.Genetics, 2(4), 280–291. 10.1038/35066065 [doi]

Heldrich, J., Sun, X., Vale-Silva, L. A., Markowitz, T. E., & Hochwagen, A. (2020). Topoisomerases Modulate the Timing of Meiotic DNA Breakage and Chromosome Morphogenesis in *Saccharomyces cerevisiae*. Genetics, 215(1), 59–73. 10.1534/genetics.120.303060 [doi]

Hillers, K. J., Jantsch, V., Martinez-Perez, E., & Yanowitz, J. L. (2017). Meiosis. WormBook: The Online Review of C. elegans Biology, 2017, 1–43. 10.1895/wormbook.1.178.1 [doi]

Hodgkin, J., Horvitz, H. R., & Brenner, S. (1979). Nondisjunction Mutants of the Nematode *Caenorhabditis elegans*. Genetics, 91(1), 67–94. 10.1093/genetics/91.1.67 [doi]

Hughes, S. E., & Hawley, R. S. (2014). Topoisomerase II is required for the proper separation of heterochromatic regions during *Drosophila melanogaster* female meiosis. PLoS Genetics, 10(10), e1004650. 10.1371/journal.pgen.1004650 [doi]

Ishiguro, T., Tanaka, K., Sakuno, T., & Watanabe, Y. (2010). Shugoshin-PP2A counteracts casein-kinase-1-dependent cleavage of Rec8 by separase. Nature Cell Biology, 12(5), 500–506. 10.1038/ncb2052 [doi]

Jaramillo-Lambert, A., Fabritius, A. S., Hansen, T. J., Smith, H. E., & Golden, A. (2016). The Identification of a Novel Mutant Allele of topoisomerase II in *Caenorhabditis elegans* Reveals a Unique Role in Chromosome Segregation During Spermatogenesis. Genetics, 204(4), 1407–1422. genetics.116.195099 [pii]

Jones, K. T. (2004). Turning it on and off: M-phase promoting factor during meiotic maturation and fertilization. Molecular Human Reproduction, 10(1), 1–5. 10.1093/molehr/gah009 [doi]

Kaitna, S., Pasierbek, P., Jantsch, M., Loidl, J., & Glotzer, M. (2002). The aurora B kinase AIR-2 regulates kinetochores during mitosis and is required for separation of homologous Chromosomes during meiosis. Current Biology : CB, 12(10), 798–812. S0960-9822(02)00820-5 [pii]

Kim, Y., Rosenberg, S. C., Kugel, C. L., Kostow, N., Rog, O., Davydov, V., Su, T. Y., Dernburg, A. F., & Corbett, K. D. (2014). The chromosome axis controls meiotic events through a hierarchical assembly of HORMA domain proteins. Developmental Cell, 31(4), 487–502. 10.1016/j.devcel.2014.09.013 [doi]

Kitajima, T. S., Sakuno, T., Ishiguro, K., Iemura, S., Natsume, T., Kawashima, S. A., & Watanabe, Y. (2006). Shugoshin collaborates with protein phosphatase 2A to protect cohesin. Nature, 441(7089), 46–52. nature04663 [pii]

Kleckner, N. (2006). Chiasma formation: chromatin/axis interplay and the role(s) of the synaptonemal complex. Chromosoma, 115(3), 175–194. 10.1007/s00412-006-0055-7 [doi]

Kleckner, N., Zickler, D., & Witz, G. (2013). Molecular biology. Chromosome capture brings it all together. Science (New York, N.Y.), 342(6161), 940–941. 10.1126/science.1247514 [doi]

Klein, F., Laroche, T., Cardenas, M. E., Hofmann, J. F., Schweizer, D., & Gasser, S. M. (1992). Localization of RAP1 and topoisomerase II in nuclei and meiotic chromosomes of yeast. The Journal of Cell Biology, 117(5), 935–948. 10.1083/jcb.117.5.935 [doi]

Krantz, I. D., McCallum, J., DeScipio, C., Kaur, M., Gillis, L. A., Yaeger, D., Jukofsky, L., Wasserman, N., Bottani, A., Morris, C. A., Nowaczyk, M. J., Toriello, H., Bamshad, M. J., Carey, J. C., Rappaport, E., Kawauchi, S., Lander, A. D., Calof, A. L., Li, H. H., . . . Jackson, L. G. (2004). Cornelia de Lange syndrome is caused by mutations in NIPBL, the human homolog of *Drosophila melanogaster* Nipped-B. Nature Genetics, 36(6), 631–635. 10.1038/ng1364 [doi]

Ladouceur, A. M., Ranjan, R., Smith, L., Fadero, T., Heppert, J., Goldstein, B., Maddox, A. S., & Maddox, P. S. (2017). CENP-A and topoisomerase-II antagonistically affect chromosome length. The Journal of Cell Biology, 216(9), 2645–2655. 10.1083/jcb.201608084 [doi]

Li, X. M., Yu, C., Wang, Z. W., Zhang, Y. L., Liu, X. M., Zhou, D., Sun, Q. Y., & Fan, H. Y. (2013). DNA topoisomerase II is dispensable for oocyte meiotic resumption but is essential for meiotic chromosome condensation and separation in mice. Biology of Reproduction, 89(5), 118. 10.1095/biolreprod.113.110692 [doi]

Liang, Z., Zickler, D., Prentiss, M., Chang, F. S., Witz, G., Maeshima, K., & Kleckner, N. (2015). Chromosomes Progress to Metaphase in Multiple Discrete Steps via Global Compaction/Expansion Cycles. Cell, 161(5), 1124–1137. S0092-8674(15)00486-9 [pii]

MacQueen, A. J., Phillips, C. M., Bhalla, N., Weiser, P., Villeneuve, A. M., & Dernburg, A. F. (2005). Chromosome sites play dual roles to establish homologous synapsis during meiosis in *C. elegans*. Cell, 123(6), 1037–1050. S0092-8674(05)01040-8 [pii]

Maeshima, K., & Laemmli, U. K. (2003). A two-step scaffolding model for mitotic chromosome assembly. Developmental Cell, 4(4), 467–480. S1534-5807(03)00092-3 [pii]

Martinez-Perez, E., Schvarzstein, M., Barroso, C., Lightfoot, J., Dernburg, A. F., & Villeneuve, A. M. (2008). Crossovers trigger a remodeling of meiotic chromosome axis composition that is linked to two-step loss of sister chromatid cohesion. Genes & Development, 22(20), 2886–2901. 10.1101/gad.1694108 [doi]

Martinez-Perez, E., & Villeneuve, A. M. (2005). HTP-1-dependent constraints coordinate homolog pairing and synapsis and promote chiasma formation during *C. elegans* meiosis. Genes & Development, 19(22), 2727–2743. 19/22/2727 [pii]

Mengoli, V., Bucciarelli, E., Lattao, R., Piergentili, R., Gatti, M., & Bonaccorsi, S. (2014). The analysis of mutant alleles of different strength reveals multiple functions of topoisomerase 2 in regulation of *Drosophila* chromosome structure. PLoS Genetics, 10(10), e1004739. 10.1371/journal.pgen.1004739 [doi]

Nasmyth, K., & Haering, C. H. (2009). Cohesin: its roles and mechanisms. Annual Review of Genetics, 43, 525–558. 10.1146/annurev-genet-102108-134233 [doi]

Nitiss, J. L. (2009). DNA topoisomerase II and its growing repertoire of biological functions. Nature Reviews.Cancer, 9(5), 327–337. 10.1038/nrc2608 [doi]

Pasierbek, P., Jantsch, M., Melcher, M., Schleiffer, A., Schweizer, D., & Loidl, J. (2001). A *Caenorhabditis elegans* cohesion protein with functions in meiotic chromosome pairing and disjunction. Genes & Development, 15(11), 1349–1360. 10.1101/gad.192701 [doi]

Petronczki, M., Siomos, M. F., & Nasmyth, K. (2003). Un menage a quatre: the molecular biology of chromosome segregation in meiosis. Cell, 112(4), 423–440. S0092867403000837 [pii]

Phillips, C. M., Wong, C., Bhalla, N., Carlton, P. M., Weiser, P., Meneely, P. M., & Dernburg, A. F. (2005). HIM-8 binds to the X chromosome pairing center and mediates chromosome-specific meiotic synapsis. Cell, 123(6), 1051–1063. S0092-8674(05)01041-X [pii]

Rankin, S. (2015). Complex elaboration: making sense of meiotic cohesin dynamics. The FEBS Journal, 282(13), 2426–2443. 10.1111/febs.13301 [doi]

Riedel, C. G., Katis, V. L., Katou, Y., Mori, S., Itoh, T., Helmhart, W., Galova, M., Petronczki, M., Gregan, J., Cetin, B., Mudrak, I., Ogris, E., Mechtler, K., Pelletier, L., Buchholz, F., Shirahige, K., & Nasmyth, K. (2006). Protein phosphatase 2A protects centromeric sister chromatid cohesion during meiosis I. Nature, 441(7089), 53–61. nature04664 [pii]

Rogers, E., Bishop, J. D., Waddle, J. A., Schumacher, J. M., & Lin, R. (2002). The aurora kinase AIR-2 functions in the release of chromosome cohesion in *Caenorhabditis elegans* meiosis. The Journal of Cell Biology, 157(2), 219–229. 10.1083/jcb.200110045 [doi]

Schindelin, J., Arganda-Carreras, I., Frise, E., Kaynig, V., Longair, M., Pietzsch, T., Preibisch, S., Rueden, C., Saalfeld, S., Schmid, B., Tinevez, J. Y., White, D. J., Hartenstein, V., Eliceiri, K., Tomancak, P., & Cardona, A. (2012). Fiji: an open- source platform for biological-image analysis. Nature Methods, 9(7), 676–682. 10.1038/nmeth.2019 [doi]

Schumacher, J. M., Golden, A., & Donovan, P. J. (1998). AIR-2: An Aurora/Ipl1-related protein kinase associated with chromosomes and midbody microtubules is required for polar body extrusion and cytokinesis in *Caenorhabditis elegans* embryos. The Journal of Cell Biology, 143(6), 1635–1646. 10.1083/jcb.143.6.1635 [doi]

Severson, A. F., Ling, L., van Zuylen, V., & Meyer, B. J. (2009). The axial element protein HTP-3 promotes cohesin loading and meiotic axis assembly in *C. elegans* to implement the meiotic program of chromosome segregation. Genes & Development, 23(15), 1763–1778. 10.1101/gad.1808809 [doi]

Severson, A. F., & Meyer, B. J. (2014). Divergent kleisin subunits of cohesin specify mechanisms to tether and release meiotic chromosomes. eLife, 3, e03467. 10.7554/eLife.03467 [doi]

Shakes, D. C., Wu, J. C., Sadler, P. L., Laprade, K., Moore, L. L., Noritake, A., & Chu, D. S. (2009). Spermatogenesis-specific features of the meiotic program in *Caenorhabditis elegans*. PLoS Genetics, 5(8), e1000611. 10.1371/journal.pgen.1000611 [doi]

Timmons, L., Court, D. L., & Fire, A. (2001). Ingestion of bacterially expressed dsRNAs can produce specific and potent genetic interference in *Caenorhabditis elegans*. Gene, 263(1-2), 103–112. S0378111900005795 [pii]

Tonkin, E. T., Wang, T. J., Lisgo, S., Bamshad, M. J., & Strachan, T. (2004). NIPBL, encoding a homolog of fungal Scc2-type sister chromatid cohesion proteins and fly Nipped-B, is mutated in Cornelia de Lange syndrome. Nature Genetics, 36(6), 636–641. 10.1038/ng1363 [doi]

Tzur, Y. B., Egydio de Carvalho, C., Nadarajan, S., Van Bostelen, I., Gu, Y., Chu, D. S., Cheeseman, I. M., & Colaiacovo, M. P. (2012). LAB-1 targets PP1 and restricts Aurora B kinase upon entrance into meiosis to promote sister chromatid cohesion. PLoS Biology, 10(8), e1001378. 10.1371/journal.pbio.1001378 [doi]

Uhlmann, F. (2016). SMC complexes: from DNA to chromosomes. Nature Reviews.Molecular Cell Biology, 17(7), 399–412. 10.1038/nrm.2016.30 [doi]

Vega, H., Waisfisz, Q., Gordillo, M., Sakai, N., Yanagihara, I., Yamada, M., van Gosliga, D., Kayserili, H., Xu, C., Ozono, K., Jabs, E. W., Inui, K., & Joenje, H. (2005). Roberts syndrome is caused by mutations in ESCO2, a human homolog of yeast ECO1 that is essential for the establishment of sister chromatid cohesion. Nature Genetics, 37(5), 468–470. ng1548 [pii]

Von Stetina, J. R., & Orr-Weaver, T. L. (2011). Developmental control of oocyte maturation and egg activation in metazoan models. Cold Spring Harbor Perspectives in Biology, 3(10), a005553. 10.1101/cshperspect.a005553 [doi]

Woglar, A., Yamaya, K., Roelens, B., Boettiger, A., Kohler, S., & Villeneuve, A. M. (2020). Quantitative cytogenetics reveals molecular stoichiometry and longitudinal organization of meiotic chromosome axes and loops. PLoS Biology, 18(8), e3000817. 10.1371/journal.pbio.3000817 [doi]

Xu, Y. X., & Manley, J. L. (2007). The prolyl isomerase Pin1 functions in mitotic chromosome condensation. Molecular Cell, 26(2), 287–300. S1097-2765(07)00190-6 [pii]

Zhang, L., Wang, S., Yin, S., Hong, S., Kim, K. P., & Kleckner, N. (2014). Topoisomerase II mediates meiotic crossover interference. Nature, 511(7511), 551–556. 10.1038/nature13442 [doi]

Zickler, D., & Kleckner, N. (1999). Meiotic chromosomes: integrating structure and function. Annual Review of Genetics, 33, 603–754. 10.1146/annurev.genet.33.1.603 [doi]

